# Sequence-dependent correlated segments in the intrinsically disordered region of ChiZ

**DOI:** 10.1101/2020.04.22.055590

**Authors:** Alan Hicks, Cristian A. Escobar, Timothy A. Cross, Huan-Xiang Zhou

## Abstract

Intrinsically disordered proteins (IDPs) account for a significant fraction of any proteome and are central to numerous cellular functions. Yet how sequences of IDPs code for their conformational dynamics is poorly understood. Here we combined NMR spectroscopy, small-angle X-ray scattering (SAXS), and molecular dynamics (MD) simulations to characterize the conformations and dynamics of ChiZ1-64. This IDP is the N-terminal fragment (residues 1-64) of the transmembrane protein ChiZ, a component of the cell division machinery in *Mycobacterium tuberculosis*. Its N-half contains most of the prolines and all of the anionic residues while the C-half most of the glycines and cationic residues. MD simulations, first validated by SAXS and secondary chemical shift data, found scant α-helices or β-strands but considerable propensity for polyproline II (PPII) torsion angles. Importantly, several blocks of residues (e.g., 11-29) emerge as “correlated segments”, identified by frequent formation of PPII stretches, salt bridges, cation-π interactions, and sidechain-backbone hydrogen bonds. NMR relaxation experiments showed non-uniform transverse relaxation rates (*R*_2_s) and nuclear Overhauser enhancements (NOEs) along the sequence (e.g., high *R*_2_s and NOEs for residues 11-14 and 23-28). MD simulations further revealed that the extent of segmental correlation is sequence-dependent: segments where internal interactions are more prevalent manifest elevated “collective” motions on the 5-10 ns timescale and suppressed local motions on the sub-ns timescale. Amide proton exchange rates provides corroboration, with residues in the most correlated segment exhibiting the highest protection factors. We propose correlated segment as a defining feature for the conformation and dynamics of IDPs.

## Introduction

Intrinsically disordered proteins (IDPs) and proteins containing intrinsically disordered regions (IDRs) comprise up to 40% of the proteomes in all life forms (1). They are involved in numerous cellular functions including regulation and signaling (2, 3). As such, dysregulation, misfolding, and aggregation of IDPs lead to many diseases (4, 5). While lacking defined tertiary structures, IDPs can exhibit conformational preferences, such as transient secondary structures and recurrent residue-residue contacts (e.g., salt bridges and cation-π interactions) (6, 7). When binding to partners, transient secondary structures may become stable (8–10), and residue-residue contacts may switch from intramolecular to intermolecular (11). Conformational dynamics may also play a particularly important role in the competition of IDPs for binding to the same partner (12) and in the binding kinetics of IDPs with partners, by dictating the binding mechanisms and binding and unbinding rate constants (11, 13–15). Clearly, the conformations and dynamics of IDPs are crucial for their cellular functions. Yet, how these properties are coded by the amino-acid sequences of IDPs is poorly understood. The present study, using an integrated experimental and computational approach, aimed to address this question for an IDR in *Mycobacterium tuberculosis* (Mtb) ChiZ, a component of the machinery responsible for cell division.

Sequence analysis and coarse-grained modeling have identified some generic determinants, in particular charged residues, for disorder and mean sizes of IDPs (16–19). In recent years, small angle x-ray/neutron scattering (SAXS/SANS) (20), fluorescence techniques including nanosecond fluorescence correlation spectroscopy (nsFCS) and single-molecule fluorescence resonance energy transfer (smFRET) (21–23), nuclear magnetic resonance (NMR) spectroscopy (24), and all-atom molecular dynamics (MD) simulations (25) have become key biophysical tools to characterize conformation ensembles of IDPs. Among these, nsFCS, NMR, and MD can also probe conformational dynamics, each with strengths on particular timescales. Scattering experiments yield information on the overall sizes and extents of disorder (20, 26). By site-specific labeling, fluorescence techniques report on the mean distances between different sites within a protein chain and the reconfiguration dynamics of the chain on the hundreds of ns timescale, as well as interactions between different protein chains (14, 27, 28).

NMR spectroscopy, based on various types of experiments, remains the only biophysical technique for characterizing both conformations and dynamics of IDPs at a residue-level resolution across timescales from picosecond (ps) to second (24). A simple telltale of intrinsic disorder is the narrow hydrogen dispersion in ^1^H-^15^N heteronuclear single quantum coherence (HSQC) spectra (29). Secondary chemical shifts, which measure the deviations from random-coil reference values, can indicate the propensities of secondary structures (30, 31). Backbone solvent exposure and interactions can be investigated using hydrogen exchange experiments for both globular and disordered proteins (32–34). Amide ^15^N spin relaxation rates report on the ps to supra-ns backbone dynamics (35). In globular proteins, relaxation data are typically analyzed through the Lipari-Szabo model-free approach, assuming separability of global tumbling motions from local backbone fluctuations (36). For disordered proteins, global and local motions are no longer separable, and interpreting relaxation data becomes a challenge. One can still model the relaxation data by fitting the NH bond vector time-correlation functions *C*_NH_(*τ*) to a sum of exponentials but the assignment of the resulting time constants to specific types of motions can be ambiguous (37–39). MD simulations can help elucidate these connections.

In recent years, it has become evident that MD force fields, traditionally parameterized for structured proteins, when applied to IDPs led to overly compact conformations (40, 41). Based on benchmarking against experimental data including SAXS profiles, FRET efficiencies, and various NMR parameters, a number of IDP-specific force fields have been proposed, including AMBER03WS/TIP4P2005 (42), various protein force fields in combination with TIP4PD water (43, 44), and CHARMM36m/TIP3Pm (45). Still, the demand for more accurate IDP force fields remains unabated, especially regarding dynamic properties. In several MD simulation studies, the *C*_NH_(*τ*) correlation functions were fitted to a sum of exponentials, but either the fitting parameters or the trajectories used for the fitting had to be adjusted in order to reach agreement with experimental data (46–51). It is thus notable that, using the AMBER14SB (52) / TIP4PD (43) force field and without adjusting *C*_NH_(*τ*), Kämpf *et al.* (53) were able to reproduce experimental relaxation data for the 26-residue N-terminal fragment of histone H4. In NMR and MD studies in which *C*_NH_(*τ*) was fitted to a sum of exponentials, 3 or 4 exponentials were typically used, and the time constants ranged from a few ps to 10 ns. While there is significant disagreement on the nature of the intermediate time scales (0.1 to 1 ns), the fastest of these motions generally is assigned to libration of the NH bond vector with respect to the peptide plane, and the slowest assigned to some form of segmental motion. One form of segmental motion arises from the simple fact that each residue is part of a polypeptide chain, which has a certain correlation length along the chain (54, 55). This type of segmental motion lacks strong sequence dependence, and can be recognized by reduced transverse relaxation rates (*R*_2_) at the chain termini (or a bell shape for the *R*_2_ vs. sequence curve), as first reported for denatured lysozyme by Schwalbe and co-workers. Although these authors also reported increases in *R*_2_ by tertiary interactions, leading to an apparent sequence dependence of *R*_2_, the latter have not received much attention in recent studies of IDPs.

The transmembrane protein ChiZ is one of a dozen or so proteins that comprise the Mtb divisome, the machinery responsible for cell division. Mtb is the causative agent of tuberculosis; its cell division has strong implications for both pathogenesis and drug resistance (56). Structural determination of divisome membrane proteins and their complexes has begun (57), but sequence analysis suggests that many of these proteins, including ChiZ, CrgA, FtsQ, FtsI, and CwsA have disordered extramembranous regions of various lengths. ChiZ consists of 165 residues (Figure 1a); the cytoplasmic N-terminal 64 residues (ChiZ1-64) are predicted to be disordered (Figure 1b,c), the next 22 residues form a transmembrane helix; and the last 53 residues form a periplasmic LysM domain that binds to peptidoglycans (58). The exact role of ChiZ in cell division is still an open question. Its full name, <u>C</u>ell wall <u>h</u>ydrolase interfering with Fts<u>Z</u> ring assembly (gene Rv2719c), may have been a misnomer, as a recent study showed that zymogram assays suggesting cell wall hydrolase activity by Chauhan *et al.* (58) likely yielded a false positive (59). On the other hand, the interference with FtsZ ring assembly remains intact. Polymerization of FtsZ (a bacterial homolog of tubulin), forming the FtsZ ring, initiates the septation step of cell division; thus correct localization of FtsZ ring is crucial for proper division (56). With increased expression of ChiZ, Mtb cells grown in macrophages were filamentous; promotion of filamentation by ChiZ overexpression in *M. smegmatis* (a non-pathogenic mycobacterium) affected the mid-cell location of FtsZ rings (58, 60). Importantly, the disordered N-terminal region and transmembrane helix sufficed for cell filamentation and FtsZ ring mislocalization (61). BATCH assays indicated that ChiZ interacts with FtsQ and FtsI, but not FtsZ, implicating an indirect mechanism for FtsZ-ring mislocalization (61).

**Figure 1.**
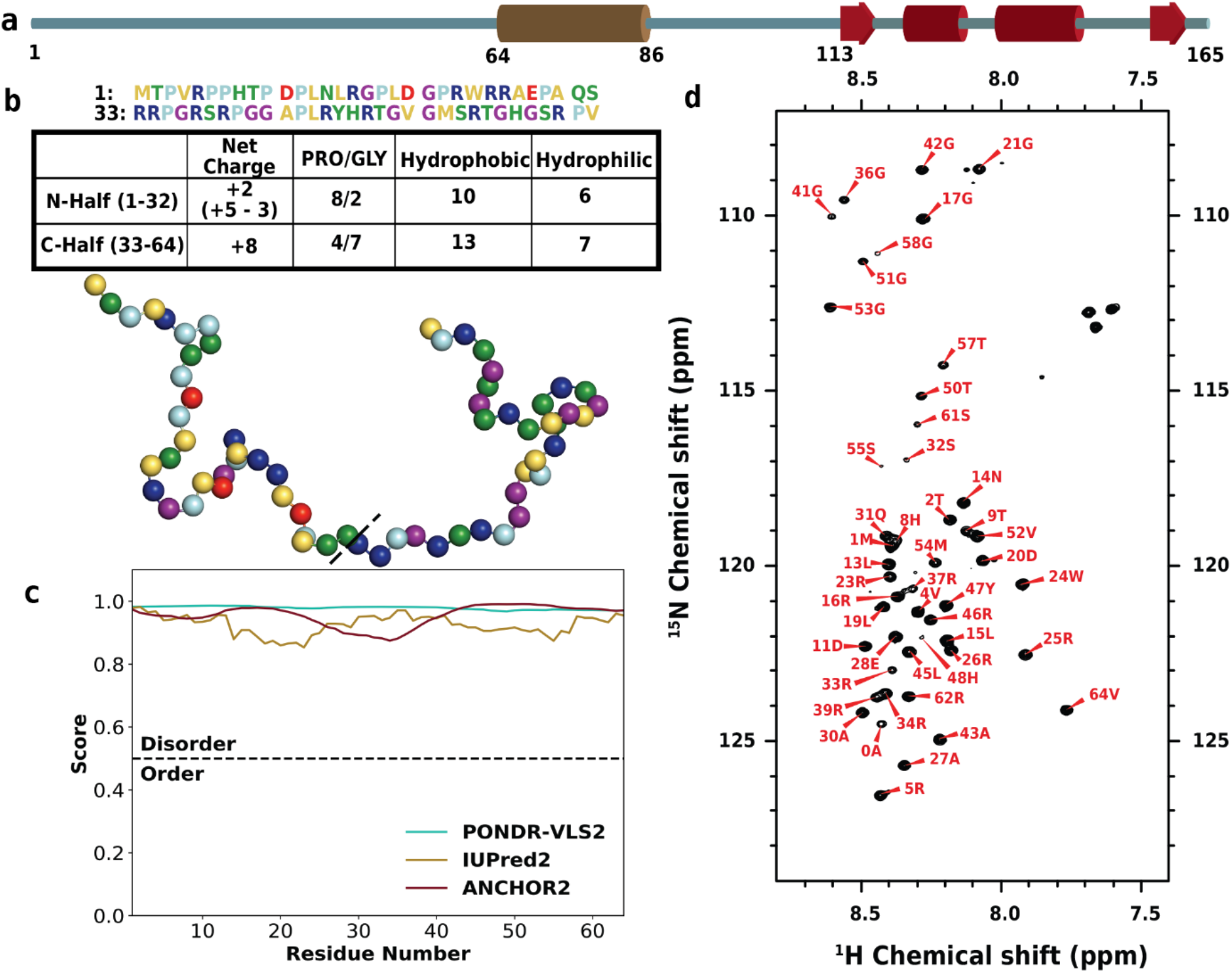
Amino-acid sequence and disorder of ChiZ. (a) Domain organization of full-length ChiZ, composed of a disordered N-terminal region (residues 1-64), a transmembrane helix, and a LysM domain (residues 113-165). (b) Sequence and amino-acid composition of ChiZ1-64. In both the sequence and an illustrative conformation, residues are colored in the following scheme: cationic, blue; anionic, red; prolines, light blue; glycines, purple; hydrophilic, green; and hydrophobic, yellow. (c) Disorder predictions from three web servers, PONDR-VLS2, IUPred2, and ANCHOR2. (d) ^1^H-^15^N HSQC spectrum, acquired on an 800 MHz spectrometer at 25 °C in 20 mM phosphate buffer (pH 7.0) with 25 mM NaCl.

Here we combined SAXS, NMR, and MD simulations to thoroughly investigate the conformational and dynamic properties of ChiZ1-64. Based on benchmarking against the SAXS profile and secondary chemical shifts, we selected the AMBER14SB/TIP4PD force field among five tested. Experimental *R*_2_ values were non-uniform along the sequence, which were recapitulated by MD simulations. The sequence-dependent dynamics can be attributed to the formation of correlated segments, stabilized by polyproline II (PPII) conformation and intra-segmental interactions. In particular, the residues with the largest amplitudes for motions on the slowest timescale (approximately 10 ns), including Asp11, Trp24, Arg25, Arg26, and Tyr47, frequently engaged, with different partners, in salt bridges and cation-π interactions. The linkage of conformation and dynamics to sequence, captured by the formation of correlated segments, will be useful for understanding IDPs and their interactions with partners.

## Results

### Sequence characteristics and disorder of ChiZ1-64

ChiZ1-64 has disparate amino-acid compositions between the N- and C-halves, in particular concerning prolines, glycines, and charged residues (Figure 1b). Of the 12 prolines (19% of the sequence), two thirds are in the N-half. In contrast, of the nine glycines (14%), seven, or nearly 80%, are in the C-half. Prolines are known to break α-helices and β-strands but promote PPII helices, whereas glycines break all secondary structures. There is a significant net charge, +10*e*, coming from 13 arginines, two aspartates, and one glutamate. All the three anionic residues are in the N-half, whereas eight, or 62%, of the cationic residues are in the C-half, resulting in the contrast between a near balance of opposite charges in the N-half and total imbalance in the C-half. Lastly we note that each half contains an aromatic residue, Trp24 in the N-half and Tyr47 in the C-half.

Giving the abundance of prolines and glycines and the high net charge, it is not surprising that the entire sequence of ChiZ1-64 is predicted to be disordered with high confidence (62–64) (Figure 1c). The ^1^H-^15^N HSQC spectrum confirms the disorder, with proton chemical shifts confined to the narrow range of 7.7 to 8.6 ppm (Figure 1d).

### SAXS profile and secondary chemical shifts

The SAXS profile, i.e., the scattering intensity *I*(*q*) as a function of *q*, the magnitude of the scattering vector (Figure 2a), especially when presented as a Kratky plot (Figure 2b), shows ChiZ1-64 as a typical IDP. The radius of gyration *R*_g_, obtained from a fit to the Debye approximation (Figure S1a), is 24.17 ± 0.05 Å. This value is slightly largely than that, 22.3 Å, predicted by a scaling relation, *R*_g_ = 2.54*N*^0.522^, deduced from a set of IDPs (65). A modest degree of expansion is also indicated by an upward tilt of the Kratky plot at high *q*.

**Figure 2.**
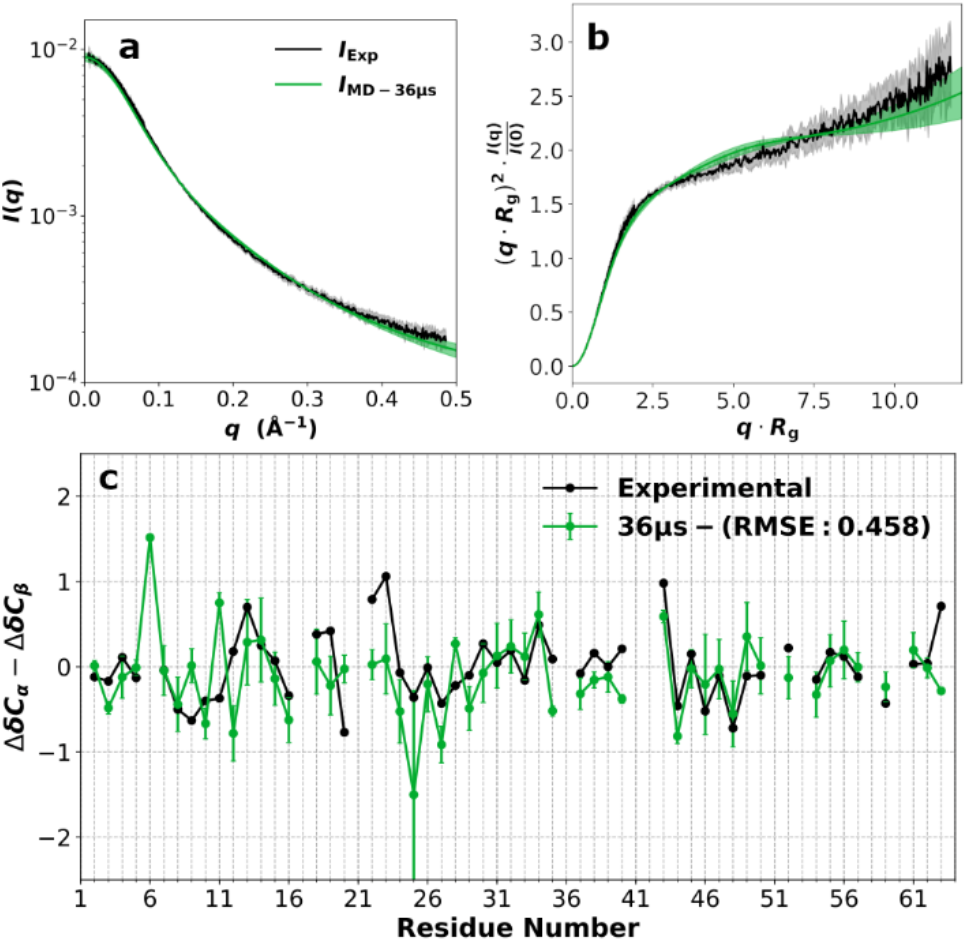
SAXS profile and secondary chemical shifts. (a) Scattering intensity *I*(*q*). (b) Kratky plot. (c) Secondary chemical shifts. Experimental data are shown in black, while the predictions from the 36-μs AMBER14SB/TIP4PD simulations are in green.

Secondary chemical shifts for C_α_ and C_β_ can indicate the presence of α-helices and β-strands (corresponding to ΔδC_α_ – ΔδC_β_ > 2 ppm and < −2 ppm, respectively) (30). For ChiZ1-64, only a single residue had |ΔδC_α_ − ΔδCβ| > 1 (Figure 2c), indicating a lack of α-helices and β-strands for the entire sequence.

### Force field validation

We used the measured SAXS profile and secondary chemical shifts to test five force fields: AMBER14SB (52) / TIP4PD (43), AMBER03WS/TIP4P2005 (42), AMBER99SB-ILDN (66) / TIP4PD (43), AMBER15IPQ/SPCEb (67), and CHARMM36m/TIP3Pm (45). In simulations totaling 2 μs, AMBER14SB/TIP4PD and AMBER03WS/TIP4P2005 did equally well and outperformed the other three force fields in matching the SAXS profile (Figure S1b-f). For secondary chemical shifts, AMBER14SB/TIP4PD was ahead of all the other four force fields (Figure S2). We further expanded the AMBER14SB/TIP4PD and AMBER03WS/TIP4P2005 simulations to 36 μs (among 12 replicate trajectories). In the expanded simulations, the AMBER14SB/TIP4PD results moved even closer toward the experimental counterparts (Figures 2, S1b, and S2a), whereas the AMBER03WS/TIP4P2005 results did not see any improvement (Figures S1c and S2b). From here on, we will focus on the AMBER14SB/TIP4PD simulations and no longer state the name of the force field.

To check the convergence of the 36-μs simulations, we calculated the *R*_g_ histograms from the 12 replicate trajectories (Figure S3). The histograms all showed broad distributions, with significant frequencies for *R*_g_ between 15 to 40 Å and mean *R*_g_ values ranging from 22.44 to 26.44 Å. Combining the 12 replicate simulations, the overall mean *R*_g_ was 24.4 Å, with a standard deviation of 1.4 Å among the replicates. This mean *R*_g_ agrees well with the experimental result. Overall, the selected force field reproduced the experimental data well for both residue-specific properties and global conformational properties.

### High PPII propensities

Consistent with the lack of α-helices and β-strands indicated by secondary chemical shifts, the contents of these secondary structures were minimal in the MD simulations (Figure 3). Two stretches of residues in the N-half, 13-15 and 30-32, formed 3_10_ helices with a moderate frequency (~7%). Note that 3_10_ helices have much lower intrinsic stability than α-helices. In addition, 3_10_ helices and anti-parallel β-sheets were formed infrequently by C-half residues (45 to 61). On the other hand, ChiZ1-64 exhibited high PPII propensities, which only the MD simulations were able to reveal. Here PPII was counted when contiguous residues (minimum of 3) fell in the PPII region on the Ramachandran map (Figure S4). Three stretches of residues sampled PPII over 50% of the time. All of them are in the N-half: residues 4-6, 10-12, and 27-29. In comparison, the highest PPII frequency in the C-half was only 36%, for residue 44. Prolines are the most direct reason for the high PPII propensities, as the high-PPII stretches in the N-half contain or border prolines at positions 3, 6, 7, 10, 12, and 29; in the C-half, residue 44 is also a proline.

**Figure 3.**
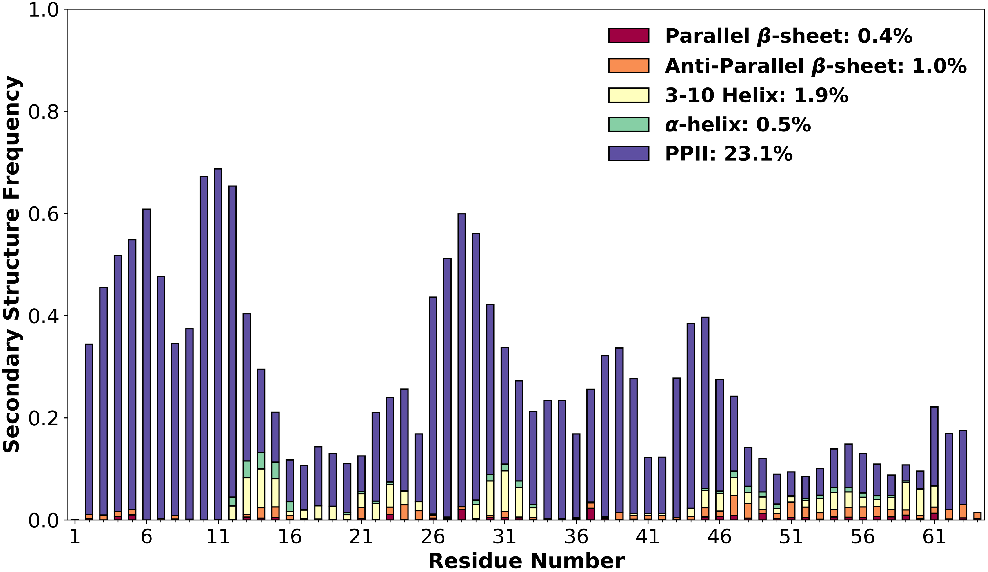
Normalized frequencies of individual residues in various types of secondary structures, according to the MD simulations.

Two of the eight N-half prolines, at positions 18 and 22, were not found in or next to high-PPII stretches, possibly because each is next to a glycine (at positions 17 and 21). Glycines may also partly explain the much lower PPII propensities of the C-half, by being next to Pro35 (at position 36), Pro40 (at positions 41 and 42), and between Pro44 and Pro63 (at positions 51, 53, 58, and 60).

Proline strongly prefers the PPII region on the Ramachandran map (Figure S4). This preference extends to the preceding residue, unless it is a glycine. However, PPII helices are only marginally stable. Unlike α-helices and β-sheets, PPII helices are not stabilized by backbone hydrogen bonds. Although prolines provide some impetus, PPII stretches may not form unless stabilized by other interactions (see below).

### Flat energy landscape in conformational space

The lack of stable secondary structures portended a high degree of diversity in the conformations sampled by ChiZ1-64. To quantify this aspect, we performed backbone dihedral principal component analysis (dPCA)(68, 69) on conformations saved from the MD simulations. Each conformation was projected onto the first two eigenmodes with the largest eigenvalues, and the distribution of the conformations in this 2-dimensional subspace was obtained. The resulting free energy surface shows a broad, shallow basin, with local barriers all less than 2 *k*_B_*T* (*k*_B_: Boltzmann constant; *T*: absolute temperature) (Figure S5a).

Another indication of the conformational diversity is provided by the closely spaced eigenvalues (Figure S6a; in a contrasting scenario where a few large eigenvalues are separated from many small eigenvalues, the former would correspond to modes involving concerted motions of a large portion of the protein, whereas the latter would correspond to localized motions). When normalized by the sum of all eigenvalues, the four largest eigenvalues were 0.029, 0.025, 0.023, and 0.021; the eigenvalues decreased smoothly with increasing mode number. The first four eigenmodes, represented by the fluctuation amplitudes of individual torsion angles, are displayed in Figure S6b-e. The amplitudes of *ϕ* angles, with the exception for those of a few glycines, were low, reflecting the fact that *ϕ* was mostly confined to the range of −50° to −150° (Figure S4). *ψ* values spanned a wide range, covering different secondary structures (−100° to 0° for α- and 3_10_ helices, and 100° to 180° for PPII helices and β-strands). Residues with high *ψ* amplitudes in the first three modes mostly were found in the two N-half stretches, 13-15 and 30-32, with a moderate 3_10_ propensity. The fourth mode mostly involved C-half residues (45 to 61) that formed 3_10_ helices and anti-parallel β-sheets infrequently.

To find a minimal set of conformations that still conveyed the overall sense of conformational diversity, we used the projections of the MD conformations in the subspace of the first two eigenmodes to group them into 16 clusters (Figure S5b) and selected one conformation from each cluster. The selection was based on a similarity score, which measured the extent of similarity of a given conformation to all other conformations in the same cluster. The highest similarity score for any conformation with all the other cluster members ranged from 0.15 to 0.19, about the same as that between two randomly chosen conformations, again highlighting the conformational diversity.

The set of 16 conformations, one from each cluster with the highest similarity score, illustrates the conformational diversity in the MD simulations (Figure 4). All these conformations contained at least one PPII stretch; five of them contained a 3_10_ helix; two contained a hybrid 3_10_-α helix (featuring both *i* to *i* + 3 and *i* to *i* + 4 hydrogen bonds); one contained an antiparallel β-sheet. Visual inspection also revealed that arginines frequently formed salt bridges with the aspartates and glutamate as well as cation-π interactions with the tryptophan and tyrosine. Furthermore, the cationic and anionic side chains frequently formed hydrogen bonds with backbone carbonyls and amides, respectively. Sometimes these interactions grew into a network. So, while the backbone conformations were diverse, salt bridges, cation-π interactions, and side chain-backbone hydrogen bonds were pervasive, albeit formed by different partners at different times.

**Figure 4.**
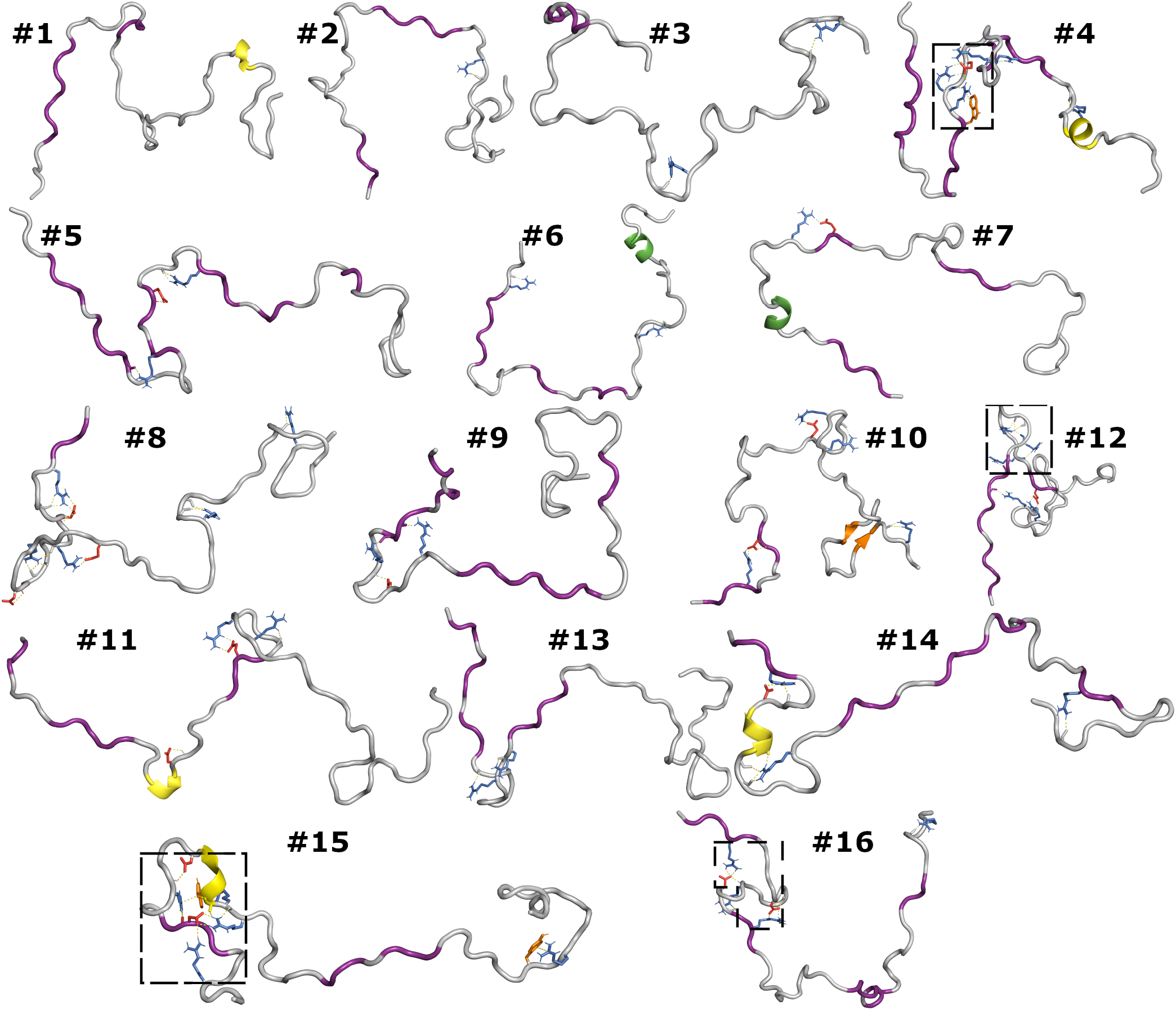
Representative conformations from the MD simulations. Secondary structures are shown by color of the backbone (PPII, 3_10_, and hybrid 3_10_-α helices in purple, yellow, and green respectively; β-sheet, orange). Cationic, anionic, and aromatic side chains involved in salt bridges, cation-π interactions, and SC-BB hydrogen bonds are shown in blue, red, and orange, respectively. Boxed regions, after enlargement, are shown in Figure 5.

### Correlated segments revealed by contact maps

To quantify these prevailing interactions formed in the MD simulations, we calculated contact frequencies between heavy atoms on any two side chains (SC-SC; Figure 5a) or between a heavy atom on any side chain and a heavy atom on the backbone of any other residue (SC-BB; Figure 5b). A contact was formed when two heavy atoms were less than 3.5 Å apart.

**Figure 5.**
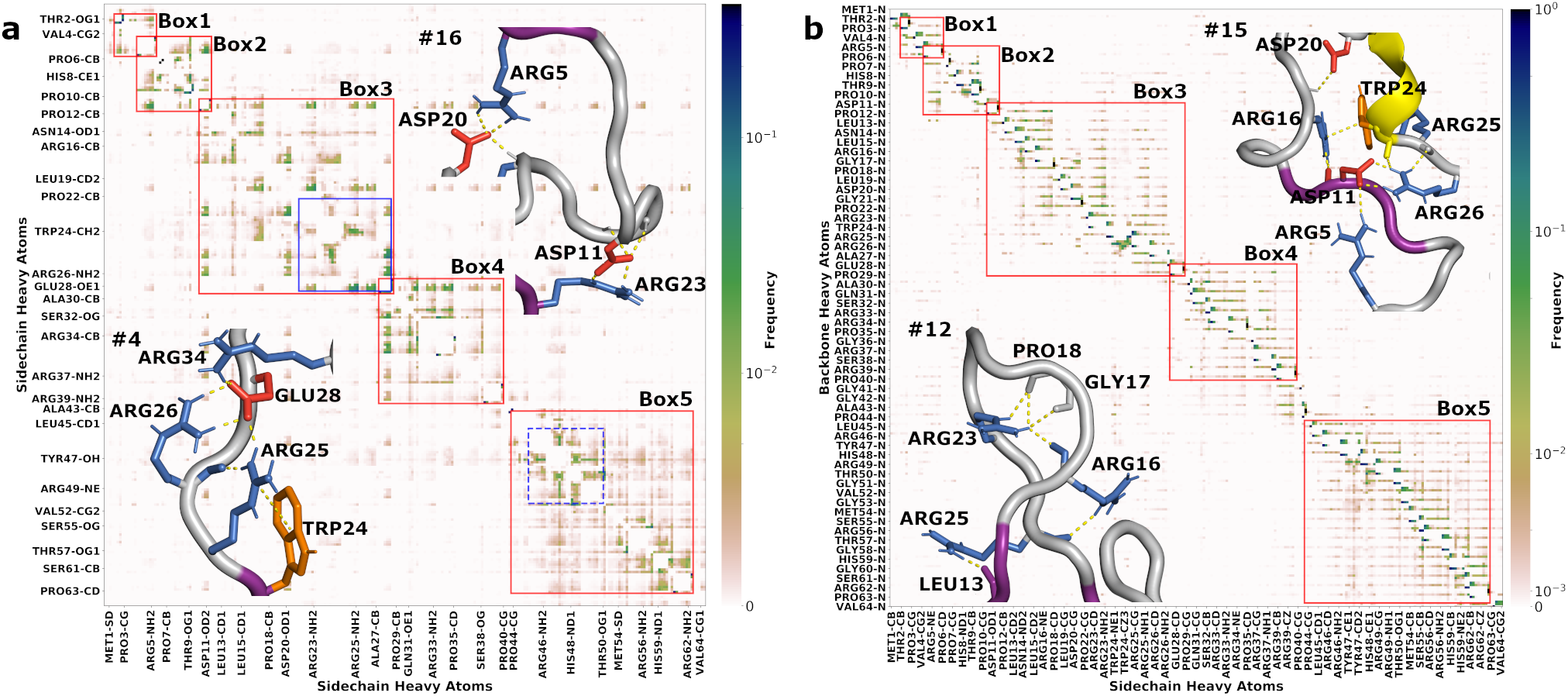
Contact maps between (a) sidechain-sidechain and (b) sidechain-backbone atom pairs. Normalized contact frequencies are shown as color gradients, which follow a linear scale for contact frequencies between 0 and 0.01 but a logarithmic scale above 0.01. Red boxes indicate segments of residues that frequently formed contacts. Blue boxes (solid, residues 23-28; dash, 46-50) highlight residues with particularly high contact frequecies. See an enlarged view of Box 3 and Box 5 in Figure S7. Enlarged views of regions from four conformations (#4, #12, #15, and #16) in Figure 4 are shown as insets to illustrate SC-SC and SC-BB interactions.

Overall, the N-half formed nonlocal SC-SC contacts much more frequently than the C-half. Residues forming SC-SC contacts with significant frequencies could roughly be grouped into five segments along the sequence (indicated by red boxes in Figure 5a). The N-half broke into three segments: Thr2 to Pro6, Arg5 to Pro12, and Asp11 to Pro29. The fourth segment, Glu28 to Pro40, straddled between the two halves. The rest of the C-half contained one more segment, Pro44 to Pro63. For several residues including Arg5, Asp11, and Glu28 (Glu28 illustrated in Figure 5a inset #4), the contacts extended beyond a single segment, explaining why every two adjacent segments in the N-half had a two-residue overlap. Contacts made by the three anionic residues, Asp11, Asp20, and Glu28, traversed the N-half (illustrated by an Asp20-Arg5 salt bridge in Figure 5a inset #16) and even extended into the entire C-half.

The most extensive interaction network was formed with Trp24, Arg25, Arg26, Glu28 at the core (Figure 5a blue solid box and inset #4; see also Figure 5b inset #15; Figure S7a). Trp24 formed cation-π interactions with Arg25 and other arginines, whereas Glu28 formed multiple salt bridges with Arg25, Arg26, and other arginines. In the C-half, the most extensive interaction network (Figure 5a blue dash box; Figure S7b) had cation-π interactions of Tyr47 with Arg46 and Arg49 at the core (e.g., Tyr47-Arg49 in Figure 4 #15). We will see that these salt bridges and π interactions align with regions of slow backbone dynamics when presenting Figures 6 and 7.

**Figure 6.**
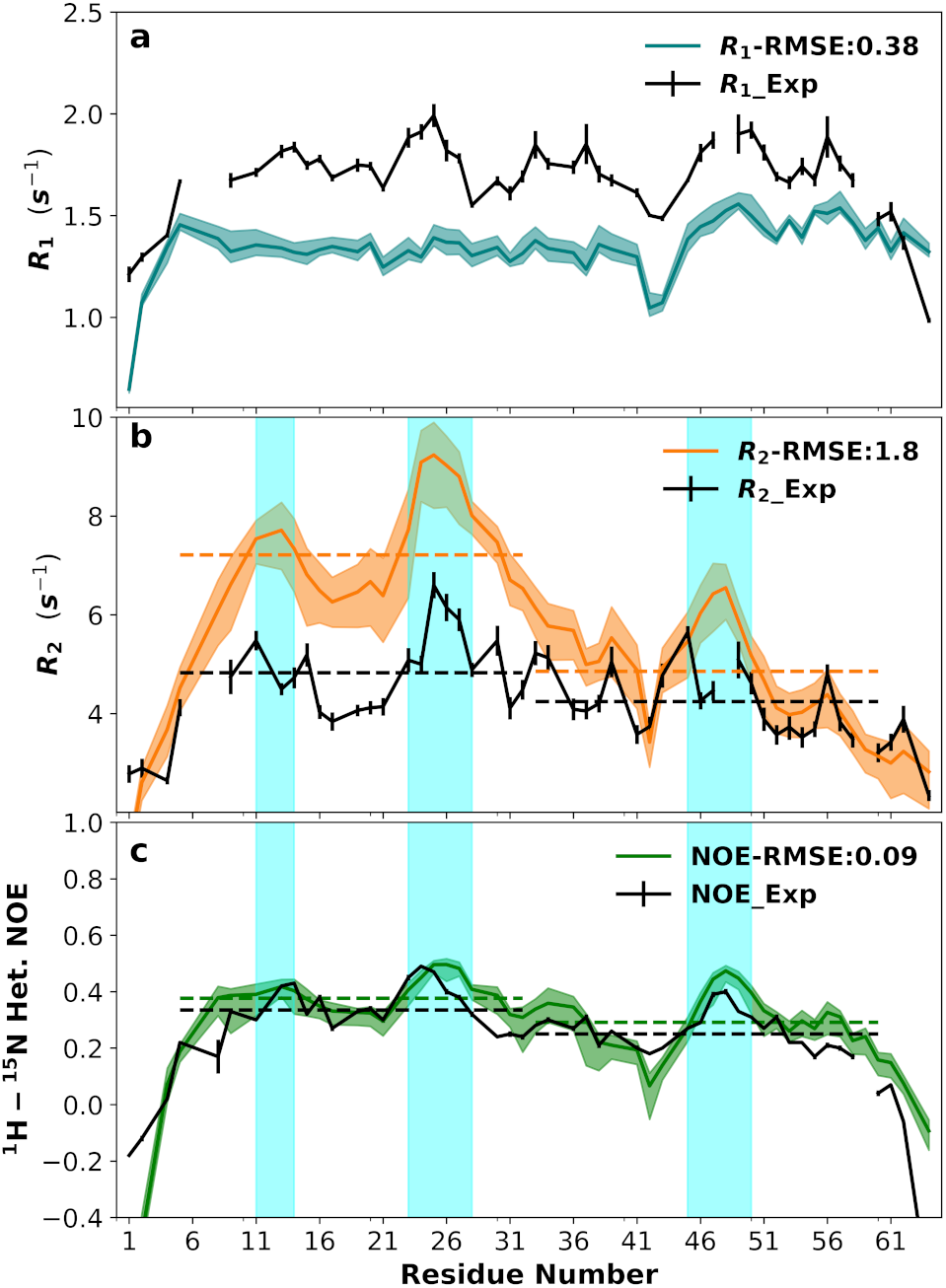
Backbone ^15^N relaxation properties. (a) *R*_1_; (b) *R*_2_; and (c) NOE. Experimental data (pH 7.0) are in black; MD results are in blue, orange, and green, with shaded bands indicating 95% confidence intervals. Dashed lines indicate averages over residues 5-32 and 33-60; shaded cyan regions highlight residues (11-14, 23-28, and 46-50) with higher-than-average *R*_2_s and NOEs.

**Figure 7.**
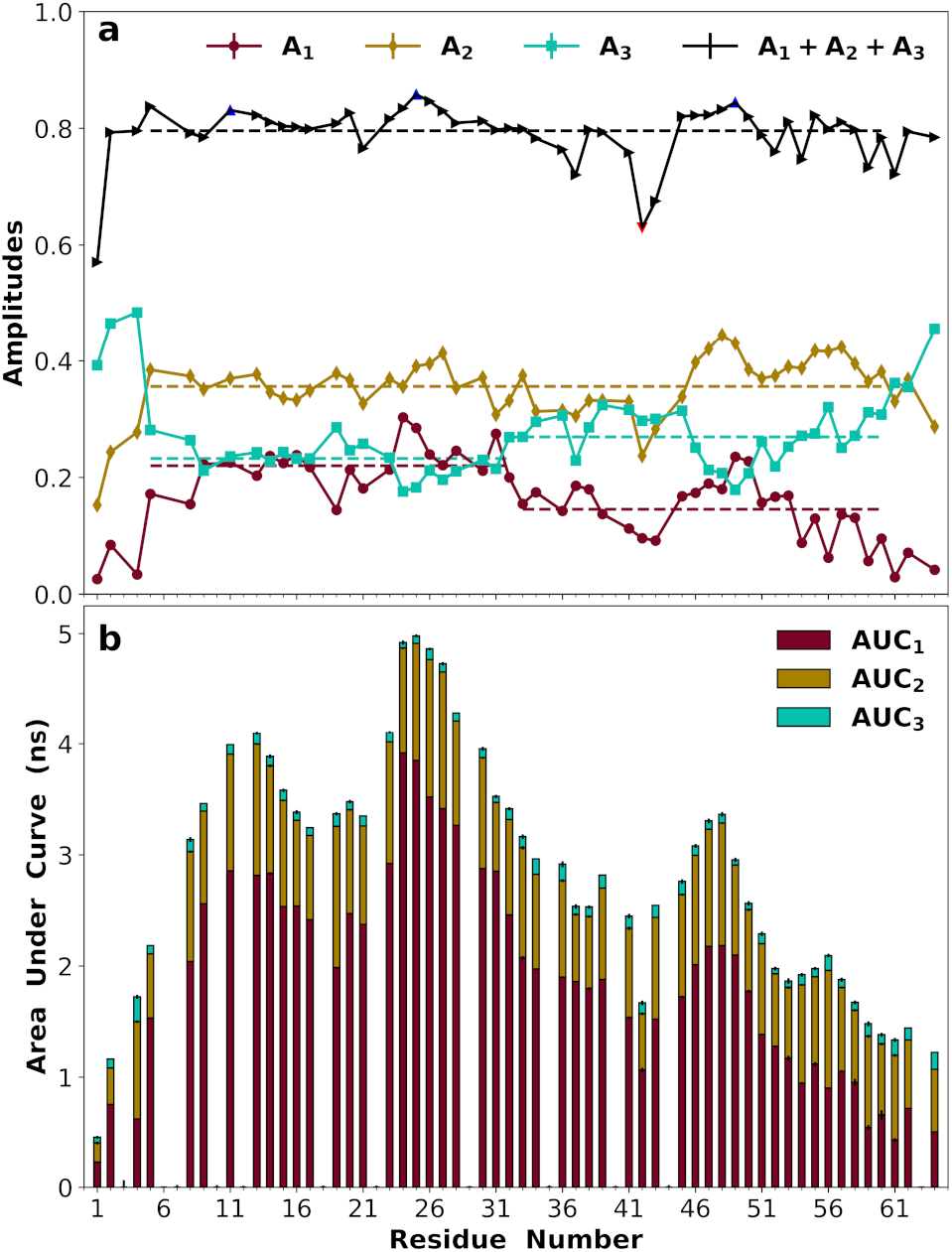
Amplitudes of backbone dynamics on three timescales. (a) Amplitudes *A*_1_, *A*_2_, *A*_3_ for exponentials with time constants in the 7-17 ns, 1.5-3.5 ns, and 0.2-0.5 ns ranges. The sum of the three amplitudes is also shown, with blue triangles (at Asp11, Arg25, and Arg49) indicating local maxmima, and red triangle (at Gly42) indicating a local minimum. Dashed lines show averages over either the entire sequence (resdiues 5-60) or the two halves (5-32 and 33-60). (b) Area under the correlation function (AUC). The contributions of the three exponentials are inidcated by bars in different colors. Fitting errors for amplitudes and propagated errors for AUC are plotted, but are smaller than the size of the symbols.

The five correlated segments each contain one or more transiently formed PPII stretches (Figure S8). The three most prevalent PPII stretches (residues 4-6, 10-12, and 27-29) identified above fall right into Boxes 1, 2, and 3. It is thus evident that SC-SC contacts contribute to the prevalence of PPII stretches. This point is clearly illustrated by the contrast between Pro29 and Pro63. These prolines are both free from the direct influence of neighboring prolines or glycines and yet differ significantly in PPII frequencies (52% for Pro29 vs. 15% for Pro63). The most likely reason for the much higher PPII frequency of Pro29 is that it is next to a stretch of residues (Trp25 to Glu28) that form extensive interactions. Pro44 is close to a stretch of residues (Arg46 to Arg48) that form less extensive interactions, and has an intermediate PPII frequency (36%).

The patterns of SC-BB contacts largely mirrored those of the SC-SC contacts. The SC-BB contacts segregated into the same five segments. Most frequent were contacts between adjacent residues, in particular hydrogen bonds between arginines and backbone carbonyls (e.g., Arg25 with the carbonyls of residues 24 and 25 as shown in Figure 5b inset #15) and between anionic residues and backbone amides. Still, nonlocal SC-BB hydrogen bonds occurred with significant frequencies in the N-half segments. For instance, Arg16 hydrogen bonded with the carbonyl of residue 25, Arg23 with residues 16, 17, and 18, and Arg25 with residue 13 as shown in Figure 5b inset #12; Arg16 with residue 11, and Asp20 with the backbone amide of residue 16 as shown in Figure 5b inset #15. There were relatively fewer nonlocal SC-BB hydrogen bonds in the C-half. All in all, the SC-BB hydrogen bonds contribute to the stability of the correlated segments identified by SC-SC contacts and, at the same time, they also directly influence backbone ^15^N relaxation and amide proton exchange rates.

### Sequence-specific backbone dynamics

In Figure 6 (black solid curves) we display the longitudinal and transverse relaxation rates (*R*_1_ and *R*_2_) and nuclear Overhauser enhancements (NOEs) of individual backbone ^15^N sites at pH 7.0. At first glance, the relaxation properties are relatively uniform across the sequence, except for the extreme four residues at each terminus, with reduced *R*_1_, *R*_2_, and NOE. The resulting “bell” shape for *R*_2_ has been suggested as arising from the residue-residue connectivity of a (denatured or disordered) polypeptide chain (54, 55). The average *R*_1_ for residues 5-60 was 1.74 s^−1^; the only pronounced deviation was a local minimum at residues Gly42 and Ala43.

Closer inspection revealed a small but systematic difference in *R*_2_ between the N- and C-halves, with mean values for residues 5-32 and 33-60 at 4.76 and 4.17 s^−1^, respectively (black dashed lines in Figure 6b). There was also a distinction in NOE between the two halves, with mean values at 0.34 for residues 5-32 and 0.25 for residues 33-60 (black dashed lines in Figure 6c). A t-test treating the N-half and C-half as two independent samples found the p-values for the differences in mean *R*_2_ and in mean NOE between the two halves to be both below 0.05 (Table S1), therefore indicating statistical significance. The overall low NOE values once again corroborate the lack of stable backbone structures. Still, the *R*_2_ and NOE data together suggest that the N-half overall has larger amplitudes for motions on the slower (e.g., 10-ns) timescale but smaller amplitudes on the faster (sub-ns) timescale than the C-half. Also worth noting are three stretches of residues, 11-14 and 23-28 in the N-half and 45-50 in the C-half (blue shading in Figure 6b,c), that had higher-than-average *R*_2_s and NOEs.

The relaxation properties at pH 4.0 showed even stronger disparity between the N- and C-halves (Figure S9). The mean *R*_2_s in the two halves were 3.10 and 2.26 s^−1^, and the mean NOEs had a wide gap, with values of 0.24 and 0.03 for the two halves. A distinction in mean *R*_1_ also became apparent between the N- and C-halves. The p-values for the differences in mean values were below 0.001 for all the three relaxation parameters, indicating strong statistical significance. A likely consequence of the decrease in pH to 4.0 is protonation of the three histidines (at positions 8, 48, and 59), which would amplify the charge imbalance in the C-half and thereby increase its disorder.

The MD simulations afforded the opportunity for a detailed interpretation of the NMR relaxation data. After evaluating the NH bond vector time-correlation functions *C*_NH_(*τ*) from the MD trajectories (at 20 ps time intervals) and fitting them to a sum of three exponentials, the resulting spectral densities were used, without any modification, to calculate the relaxation properties. The results were close to the experimental counterparts, but with systematic underestimates in *R*_1_ and overestimates in *R*_2_ (colored solid curves). The root mean square errors (RMSEs) relative to the experimental data (excluding the extreme residue at each terminus) were 0.38 s^−1^ for *R*_1_, 1.8 s^−1^ for *R*_2_, and 0.09 for NOE. Importantly, the MD simulations recapitulated the sequence-dependent features of the experimental data, including: (1) the overall differences in *R*_2_ and NOE between the N- and C-halves (as indicated by disparate mean values in the two halves, shown as colored dashed lines in Figure 6b,c); (2) the three stretches of residues showing local maxima in *R*_2_ and NOE (blue shading in Figure 6b,c); and (3) the local minimum in *R*_1_ at residues 42 and 43 (Figure 6a).

### Amplitudes of backbone dynamics on different timescales

Given the above qualitative agreement with the experimental data, we now report on the MD results for *C*_NH_(*τ*), specifically their tri-exponential fits (with time constants *τ*_1_, *τ*_2_, and *τ*_3_, ordered from large to small, and amplitudes *A*_1_, *A*_2_, and *A*_3_; see Figure S10 for representative fits). It is important to note that we did not restrain the sum of the amplitudes, *A*_sum_ = *A*_1_ + *A*_2_ + *A*_3_, to be 1. Implicitly, we assumed that the missing amplitude, 1 − *A*_sum_, represented an ultrafast decay that occurred before the first time point, 20 ps, at which we evaluated *C*_NH_(*τ*). Indeed, adding an ultrafast decay component with an amplitude of 1 − *A*_sum_ and a time constant of τ_f_ = 10 ps largely made up for some underestimates of the tri-exponential fits at short times (Figure S10 insets). The mean ± standard deviation of *A*_sum_ for non-terminal residues were 0.80 ± 0.04 (black dashed line in Figure 7a). In comparison, the order parameters for NH libration calculated after superimposing the peptide plane were 0.933 ± 0.001, implicating additional contributions (e.g., rapid fluctuations of the *ϕ* and *ψ* angles adjoining the peptide plane (53)) to the ultrafast decay. Interestingly, three local maxima in *A*_sum_, at Asp11, Arg25, and Arg49, apparently coincided with the residues showing higher-than-average *R*_2_s and NOEs (triangles filled in blue in Figure 7a), whereas the minimum in *A*_sum_ at Gly42 (triangles filled in red in Figure 7a) coincided with the residues showing lower-than-average *R*_1_s.

The mean ± standard deviation of the three time constants were 11.5 ± 2.4, 2.4 ± 0.5, and 0.34 ± 0.06 ns for the non-terminal residues. The three exponentials with these time constants each contribute most to a different relaxation property, specifically, with the slow, intermediate, and fast timescales controlling *R*_2_, *R*_1_, and NOE, respectively. The amplitudes (*A*_2_) associated with the intermediate time constant were nearly uniform along the sequence (at 0.36 ± 0.04), except for two very low values, at Gly42 and Ala43 (Figure 7a). These results on *A*_2_ largely explain the corresponding behavior of *R*_1_ presented above, i.e., near constancy except for higher-than-average values for Gly42 and Ala43. On the other hand, *A*_1_ and *A*_3_ showed disparities between the N- and C-halves. In the N-half, the *A*_1_ and *A*_3_ averages were nearly the same, at 0.22 and 0.23, respectively. In the C-half, the *A*_1_ average moved down to 0.15 while the *A*_3_ average moved up to 0.27. Given the near constancy of *A*_2_ and *A*_sum_, the opposite movements of *A*_1_ and *A*_3_ were inevitable. The disparity in *A*_1_ between the two halves explains the corresponding disparity in *R*_2_, with lower *A*_1_ values in the C-half accounting for the longer *R*_2_s (i.e., weaker transverse relaxation) in that half. Likewise, the disparity in *A*_3_ between the two halves explains the corresponding disparity in NOE, with higher *A*_3_ values in the C-half accounting for the lower NOEs (i.e., higher flexibility) in that half.

The area under the *C*_NH_(*τ*) curve (AUC) equals the spectral density, *J*(0), at zero frequency, to which *R*_2_ is particularly sensitive. The AUC values (and their contributions from the three exponentials) are displayed in Figure 7b. Two patterns are apparent (which are also true of the *A*_1_ component). First, the N-half overall had higher AUCs than the C-half (with averages at 3.8 and 2.4 ns, respectively). Second, there were three local maxima, at residues 11-13, 24-27, and 47-48. These were the same maxima as identified based on *A*_sum_ (Figure 7a), but were now much more conspicuous. They explain the higher-than-average *R*_2_s of the involved residues (Figure 6b).

Ultimately, the higher amplitudes (*A*_1_) for the slow timescale (and higher AUCs) of the N-half came from the more frequent SC-SC and SC-BB contacts in this half, in particular salt bridges, cation-π interactions, and SC-BB hydrogen bonds mediated by charged residues, resulting in correlated segmental motions. Indeed, the two most extensive interaction networks, centered around residues 24-27 and 46-49, were directly responsible for the local maxima in AUC and the corresponding local maxima in *R*_2_ at these residues. In contrast, for Gly41 and Gly42, the absence of a sidechain not only allows them to access the left handed side of the Ramachandran map (Figure S4), but also precludes them from forming any SC-SC or SC-BB contacts (Figure 5), resulting in much faster backbone dynamics.

### Non-uniform amide proton exchange rates along the sequence

Amide proton exchange rates (*k*_ex_; Figure 8a) further corroborated the presence of correlated segments suggested by the NMR relaxation experiments and MD simulations. The average *k*_ex_ of the N-half, 3.9 s^−1^, was less than one third of the counterpart of the C-half, 12.4 s^−1^.

**Figure 8.**
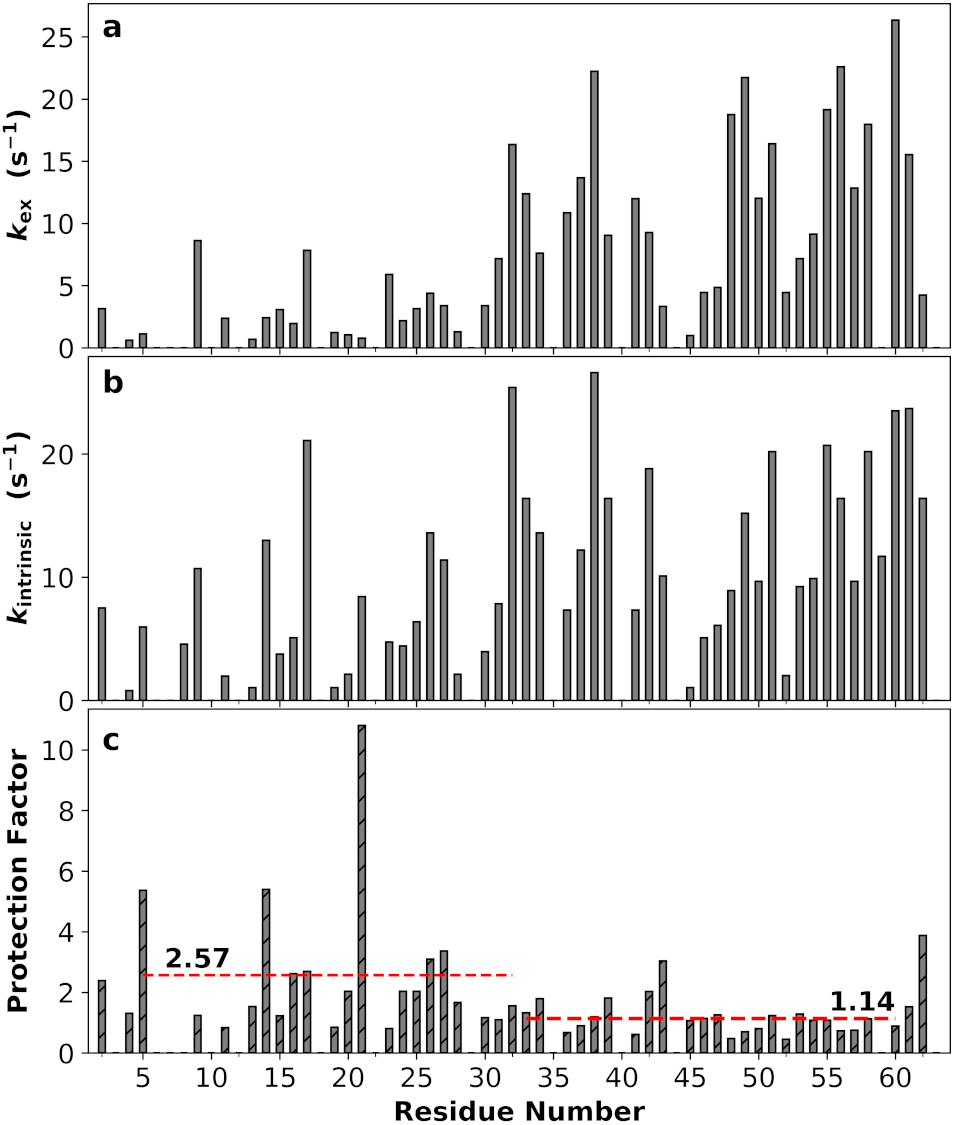
Backbone amide proton exchange rates (*k*_ex_; pH 7.0), intrinsic exchange rates (*k*_intrinsic_), and protection factors. (a) *k*_ex_. (b) *k*_intrinsic_. (c) Protection factors. The mean protection factors for residues 5-32 and 33-60 are shown by red dashes.

*k*_ex_ is strongly dependent on the amino-acid sequence. We calculated the intrinsic exchange rates (*k*_intrinsic_) from the sequence using the SPHERE server (Figure 8b) (70). By taking the ratio *k*_intrinsic_/*k*_ex_, we obtained the protection factors (Fig. 8c). Interestingly, for the C-half residues, *k*_ex_ values were all close to *k*_intrinsic_ values, indicating there is very little influence beyond the immediate amino-acid sequence (average protection fact at 1.14). In contrast, for most of the N-half residues, the protection factors were higher than 1, averaging at 2.57. A t-test showed that the difference in mean protection factor between the N- and C-halves is statistically significant (p-value at 0.015; Table S1). This difference can be nicely explained by the disparities in SC-SC and SC-BB contacts between the two halves. In particular, the two residues with the highest protection factors (residue 14 at 10.8 and residue 21 at 5.4) are located in the most correlated segment (residues 11-29, identified by extensive interactions and high *R*_2_s and NOEs).

## Discussion

By combining NMR, SAXS, and MD simulations, we have characterized the conformations and dynamics of ChiZ1-64 and delineated their linkage to the amino-acid sequence. The conformations of ChiZ1-64 were diverse, with the only notable feature being high propensities of PPII stretches, especially in the N-half. Backbone ^15^N relaxation experiments revealed non-uniform *R*_2_s and NOEs along the sequence, with high values for residues 11-14 and 23-28. These or neighboring residues also have high protections factors for amide proton exchange. MD simulations recapitulated these observations and suggest that the reason for the non-uniform dynamics is the formation of correlated segments, which are stabilized by PPII stretches, salt bridges, cation-π interactions, and sidechain-backbone hydrogen bonds. Moreover, the extent of segmental correlation is sequence-dependent: segments where internal interactions are more prevalent manifest elevated “collective” motions and suppressed local motions.

Similar to ChiZ1-64, sequence-specific backbone dynamics has been reported on a number of other IDPs using NMR, some in combination with MD simulations (38, 48–50, 53, 55, 71, 72). Whereas stable secondary structures such as α-helices and β-hairpins can certainly lead to slow backbone dynamics (38, 48, 50, 53), as demonstrated here for ChiZ1-64, interaction networks, in particular those mediated by charged and aromatic residues, can lead to the formation of correlated segments, which can have slow dynamics even when the backbone remains disordered. We propose correlated segment as a defining feature for the conformation and dynamics of IDPs. Contact maps provide a way to identify correlated segments and characterize their stabilizing interactions. For example, it is interesting to investigate whether cation-π or other types of interactions contribute to the slow dynamics of two tryptophans in the C-terminal domain of the nucleoprotein of Sendai virus (38, 48), or the precise interactions that may be responsible for the slow dynamics of two stretches of residues in HOX transcription factors (72). Accumulation of such knowledge over a large number of IDPs will advance our understanding of how amino-acid sequences of IDPs, through the formation of correlated segments, code for dynamics.

There is pressing need for continued development of IDP force fields. The AMBER14SB/TIP4PD force field selected here based on benchmarking against SAXS profile and chemical shifts also performed reasonably well for dynamic properties. Coincidentally, the same force field was also selected by Kämpf *et al.* (53) from comparison with backbone ^15^N relaxation data. Still, for ChiZ1-64, the MD results had an apparent systematic underestimation of *R*_1_ and overestimation of *R*_2_. The opposite deviations suggest an exaggeration of the longest timescale in the NH bond vector time-correlation functions. To test this idea, we scaled down the three time constants from the tri-exponential fits by a factor 1 + τ_i_/τ_s_, with τ_s_ on the order of 10 ns, along with the addition of an ultrafast decay component with time constant τ_f_ = 10 ps and amplitude 1 − *A*_sum_ noted above. With τ_s_ = 16.75 ns, the systematic errors were reduced for *R*_1_ and almost eliminated for *R*_2_; the NOE calculations maintained the good agreement with the experimental data (Figure S11). Whether TIP4PD indeed makes ns dynamics too slow and, if so, how to improve this promising water model warrant further studies. It is also possible the Langevin thermostat affects the dynamics of ChiZ1-64. Dynamic properties of IDPs have much to contribute in force field validation and improvements.

Although the functional role of ChiZ in Mtb cell division remains open, it may involve interactions with other divisome proteins including FtsQ and FtsI (61). Like ChiZ, both of the latter proteins contain disordered cytoplasmic regions high in charged amino acids. The interactions between all these IDRs may lead to fuzzy complexes. Moreover, these highly charged IDRs are also very likely to associate with the highly anionic Mtb membranes. The conformational and dynamic characterization of the ChiZ IDR in isolation done here will set the stage for studying these more complex systems. Given the disparity between the two halves of the ChiZ IDR, we expect the N-half to be more calcitrant while the C-half more adaptive in interacting with the various partners.

### Experimental and computational methods

#### Protein expression and purification

Expression of ChiZ1-64 containing an N-terminal His-tag with a TEV protease cleavage site was performed in *E. coli* BL21 cells. Cells were grown at 37 °C until O.D. at 600 nm was 0.7 and then 0.4 mM IPTG was added to induce expression for 5 hours at 37 °C. For ^13^C-^15^N uniformly labeled samples, protein expression was performed in M9 media containing 1 g of ^15^N ammonium chloride and 2 g of ^13^C labeled glucose.

ChiZ1-64 was initially purified using nickel affinity chromatography. Cells were resuspended in a lysis buffer (20 mM tris-HCl pH 8.0 containing 500 mM NaCl and 6 M urea) and lysed by a French Press. The lysate was centrifuged at 12,000 × g for 40 min to remove insoluble material. After that, the lysate was loaded onto a Ni-NTA resin column (Qiagen). Th column was washed with a washing buffer (20 mM tris-HCl pH 8.0 containing 500 mM NaCl and 60 mM imidazole) and then eluted with 400 mM imidazole.

After nickel affinity chromatography, the fractions containing the protein were pooled and treated with TEV protease. His-tag and TEV protease were removed by passing the protein sample through a Ni-NTA column. Further purification proceeded by cation exchange chromatography. ChiZ1-64 was dialyzed against the NMR buffer (20 mM sodium phosphate at pH 7.0 plus 25 mM NaCl) and then loaded into an SP column (GE Healthcare). The column was washed with the NMR buffer containing 200 mM NaCl, and eluted with 500 mM NaCl. Fractions containing the protein were concentrated and dialyzed against the NMR buffer for further experiments.

#### Small angle X-ray scattering

SAXS experiments were performed on the DND-CAT 5ID-D beam line at the Advanced Photon Source of the Argonne National Laboratory. The X-ray wavelength was 1.2398 Å. X-ray scattering intensities were collected using Rayonix LX170HS CCD detectors positioned at 200.92 mm (0.0014 < *q* < 0.08 Å^−1^) and 1014.2 mm (0.077 < *q* < 0.485 Å^−1^). ChiZ1-64 was at 9.1 mg/mL in the NMR buffer. X-ray exposure time was limited to 5 seconds to minimize protein degradation. Data processing was done using the ATSAS software package (73).

### NMR spectroscopy

Samples for solution NMR experiments were prepared in the NMR buffer containing 10% D_2_O and 50 μM DSS (*2*,*2-dimethyl-2-silapentane-5-sulfonic acid, for referencing).* All experiments were performed at 25 °C in an 800 MHz magnet equipped with cryoprobe. Sequential backbone assignment was performed using standard HNCO, HN(Cα)CO, HNCαCβ, and CαCβ(CO)NH experiments. Data were processed using Topspin 2.1 (Bruker), and analyzed using the CCPNmr software.

Backbone dynamics was characterized by measuring amide ^15^N *R*_1_ and *R*_2_ relaxation times. *R*_1_ measurements were performed using the Bruker pulse sequence hsqct1etf3gpsi while *R*_2_ relaxation times were measured using the Carr-Purcell-Meiboom-Gill experiment (hsqct2etf3gpsi). Relaxation delays used between scans were 6 s for *R*_1_ and 4 s for *R*_2_. In addition, using the Bruker pulse sequence hsqcnoef3gpsi, the ^1^H-^15^N heteronuclear NOE value for each residue was obtained as the ratio of signal intensities collected with and without proton saturation, with a 10-s relaxation delay between scans. The same settings were used for relaxation measurements at pH 4.0.

Measurement of amide proton exchange rates was carried out using the CLEANEX-PM pulse sequence (fhsqccxf3gpph). CLEANEX-PM spin lock times were 10, 15, 20, 25, 30, 40, 50, 75, and 100 ms. A fast HSQC reference spectrum was collected using the same pulse sequence parameters. All experiments were run with a relaxation delay of 3 s. Amide proton exchange rates were calculated by fitting equation 1 in Hwang *et al.* (74) to the signal intensities for each residue at different spin lock times. Intrinsic exchange rates were calculated using the SPHERE server (https://protocol.fccc.edu/research/labs/roder/sphere/sphere.html) (70). Protection factors were calculated as *k*_*intrinsic*_/*k*_*ex*_ (75).

Significance of the differences in the means of *R*_1_, *R*_2_, NOE, and protection factor between the two halves of ChiZ1-64 was analyzed by the independent-samples t-test assuming unequal variances (Welch’s t-test), using the *scipy.stats* module in python.

### Molecular dynamics simulations

Five force fields were tested on ChiZ1-64 in solution: AMBER14SB (52) / TIP4PD (43) (FF14D), AMBER03WS/TIP4P2005 (42) (FF03WS), AMBER99SB-ILDN (66) / TIP4PD (43) (FF99D), AMBER15IPQ/SPCEb (67) (FF15IPQ), and CHARMM36m/TIP3Pm (45) (C36M). MD simulations of ChiZ1-64 (with ACE and NME caps), started from an extended conformation generated using tleap in AmberTools16 (76), were performed in AMBER16 (76) (and extended in AMBER18 (77)). The AMBER topology file of the TIP4PD water model was from https://github.com/ajoshpratt/amber16-tip4pd. The FF14D, FF99D, and FF15IPQ simulation systems were set up using tleap, in an orthorhombic box with 12 Å of space to all sides of the protein. 25 mM NaCl was added to the solution along with 10 neutralizing Cl^−^ ions. The FF03WS system was built in GROMACS 2016.4(78, 79) to match the AMBER systems and then converted to AMBER format using GROMBER in PARMED (76). The C36m system was built using the solution builder in CHARMM-GUI (80), again to match the AMBER systems, and exported to AMBER format using CHAMBER (81). The total numbers of atoms in the systems ranged from 112,000 to 162,000.

To start, energy minimization was carried out using *sander*, for 2000 steepest descent steps, followed by 3000 conjugate gradient steps. Subsequently, temperature equilibration, pressure equilibration, and production run were done using *pmemd.cuda* on GPUs (82). Under constant volume, the temperature was ramped from 0 K to 300 K in 40 ps and then maintained at 300 K for 60 ps, using the Langevin thermostat with a friction coefficient of 3 ps^−1^, at a 1 fs timestep. The simulations then switched to constant pressure (Berendsen thermostat at 1 atm, with pressure relaxation time of 2 ps), and the timestep increased to 2 fs. The first 3 ns was nominally pressure equilibration and the remaining simulations counted as the production run. The non-bonded cutoff was 10 Å in all simulations, except in C36m where a force switch was imposed between 10 and 12 Å. All bonds connected to hydrogens were constrained using the SHAKE algorithm (83).

For each of the force fields tested, 4 replicate simulations started with different random seeds were run for 500 ns each. Simulations in two of the force fields, FF14D and FF03WS, were expanded to 12 replicates, each lasting 3 μs. Snapshots were saved every 20 ps for analysis. In only 1.7% of snapshots ChiZ1-64 came within the non-bonded cutoff (10 Å) from its periodic images.

### Calculation of SAXS profiles

From MD conformations, SAXS profiles were calculated using the FOXS code (84). The optimal parameter for the hydration shell scattering density was selected for each water model according to Henriques *et al.* (85). For each trajectory, a SAXS profile was calculated on every 10th saved conformation and then an average was taken over these conformations. The resulting SAXS profile was linearly scaled to best match the experimental counterpart. The average of the scaled SAXS profiles over replicate simulations was taken as the MD prediction. In cases where the number of replicates is 12, we also report the standard deviation among the replicates at each *q* as a measure of the calculation error.

All graphs were plotted using *matplotlib* and *seaborn* in python3.

### Calculation of chemical shifts

Chemical shifts were calculated using the SHIFTX2 code (86) (at 300 K and pH 7). For reach residue, the corresponding random-coil chemical shifts, generated from the ncIDP database (87), were subtracted to yield Cα and Cβ secondary chemical shifts (without any scaling). Details for averages and standard deviations largely followed the protocol for SAXS profiles.

### Radius of gyration, secondary structures, and hydrogen bonds

*cpptraj* was used for determining the radius of gyration, secondary structures (by implementing DSSP (88)), and hydrogen bonds. DSSP was modified to include PPII, following Mansiaux *et al.* (89). Specifically, three or more consecutive residues that were classified as coil and fell into the PPII region of the Ramachandran map were reclassified as PPII.

### Dihedral principal component analysis

Dihedral principal component analysis (dPCA)(68, 69) was done through *cpptraj*, yielding 248 eigenmodes for the backbone *ϕ* and *ψ* angles of the 62 non-terminal residues of ChiZ1-64 (*ϕ* and *ψ* each was represented by its sine and cosine). To display the energy landscape in conformational space, the histogram of the projections of the saved snapshots along the two dPCA eigenmodes with the largest eigenvalues was calculated, and then converted to a free energy surface according to the Boltzmann relation. These two projected coordinates were also used to group the snapshots into 16 clusters using the hierarchical Ward agglomerative algorithm. The snapshot that had the highest similarity score to all the members in a cluster was selected as the representative. The similarity score was defined as (90)

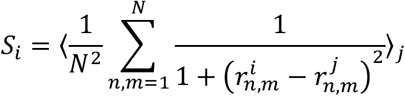

where 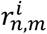 denotes the distance between atoms *n* and *m* in snapshot *i*, *N* is the total number of atoms in ChiZ1-64, and the average over *j* ran over all the snapshots in the given cluster.

The contribution from the fluctuations of torsion angle *n* to eigenmode *k* was determined by the amplitude of this eigenmode’s components, 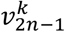 and 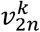, for the sine and cosine of the torsion angle (denoted by indices 2*n* – 1 and 2*n*). Specifically, this contribution was 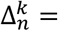 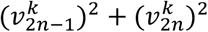 (69). Note that the sum of 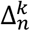 over all the torsion angles is 1.

### Contact maps

*mdtraj* (91) was used to load the trajectories, select atoms, and calculate distances between sidechain and sidechain or sidechain and backbone heavy atoms, excluding pairs from the same residue. Two heavy atoms were considered in contact if they were within 3.5 Å of each other. For each pair, the fraction of snapshots in which contacts formed was recorded.

Contacts formed by the two aromatic residues, Trp24 and Tyr47, with arginines were further examined to see whether they were cation-π interactions. The distance between the centers of mass of the indole or phenol ring and of the cationic group (including N_ε_, C_ζ_, N_η1_, and N_η2_), and the angle between the line connecting these two points and the normal of the ring were calculated. The overwhelming majority of Trp24 contacts with Arg16, Arg25, and Arg26 had the above distance < 5 Å and the above angle < 60°, and hence were deemed cation-π interactions. The same was true of Tyr47 contacts with Arg46 and Arg49.

### NMR relaxation properties

From each trajectory, the NH bond vector time-correlation function for each non-proline residue was calculated as a time average: *C*_NH_(τ) = ⟨*P*_2_[*n*(*t* + τ) · *n*(*t*)]⟩_*t*_, where *P*_2_(*x*) is the second-order Legendre polynomial and ***n***(*t*) is the NH bond unit vector at time *t*. Each correlation function, in the τ range from 20 ps and 25 ns, was then least-squares fitted to a sum of exponentials,

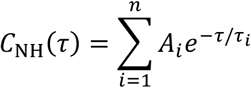

using the Levenberg-Marquardt algorithm from the *scipy.optimize.curve_fit* module in python. Note that the sum of the amplitudes was not restricted to 1, in contrast to most other studies, under the assumption that an ultrafast decay was completed by τ = 20 ps, thereby accounting for some missing amplitudes (see below). To determine the optimal number of exponentials for modeling the simulation data, we compared the chi-squares (*χ*^2^) of the fits with increasing *n*, starting at *n* = 2. Any fit with fitting errors higher than 10% of any fitted parameters was rejected. An increase from *n* exponentials to *n* + 1 exponentials was accepted only if 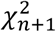 was substantially smaller than 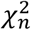, specifically if

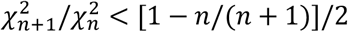

This procedure led to *n* = 3 as the optimum for all residues. The three time constants were ordered as τ_1_ > τ_2_ > τ_3_. When displaying fitting parameters (Figure 7), the fitting errors from the tri-exponential fit are also shown.

After the tri-exponential fit, the spectral density was obtained as

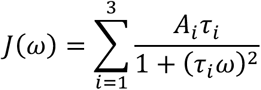

Finally the longitudinal and transverse relaxation times and the NOE were obtained as (35)

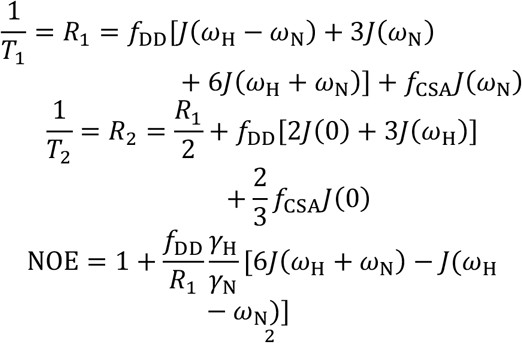

Here 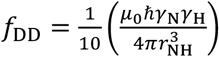 and 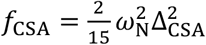 represent contributions to ^15^N spin relaxation by N-H dipole-dipole interactions and the nitrogen chemical shift anisotropy, respectively. The meanings of the other symbols are: *μ*_0_, permittivity of free space; 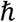, reduced Plank constant; γ_N_ and γ_H_, gyromagnetic ratios of nitrogen and hydrogen; *ω*_H_ = γ_H_*B*_0_, Larmor frequency of hydrogen (800 MHz in our case); *ω*_N_, counterpart of nitrogen; *r*_NH_, N-H bond length (set at 1.02 Å); and ∆_CSA_ (= −170 ppm), chemical shift anisotropy of nitrogen. For each relaxation property, we report the discrepancies between the measured and predicted values as the root mean squared error (RMSE), calculated over the entire ChiZ1-64 sequence but excluding the first and last residues. A bootstrapped 95% confidence interval was obtained to determine the error in the calculated *R*_1_, *R*_2_ and NOE.

We also considered two modifications of the tri-exponential NH bond vector time-correlation functions. The first was to include an ultrafast decay component, with time constant τ_f_ (< 20 ps) and an amplitude of 1 – *A*_sum_. The second was to account for the possibility that the longest timescale was exaggerated by the AMBER14SB/TIP4PD force field selected here. We hence tested scaling down the three time constants from the tri-exponential fits by a factor 1 + τ_*i*_/τ_s_, with τ_s_ of the order of 10 ns. This scaling has little effect on τ_2_ and τ_3_, but reduced the longest time constant τ_1_ by roughly half, from 7-17 ns to 5-8 ns.

### Data availability

The chemical shifts of ChiZ1-64 have been deposited in BMRB (accession # 50115). Python scripts written for the NMR relaxation analysis are available on GitHub at https://github.com/achicks15/CorrFunction_NMRRelaxation.

## Acknowledgments

This work was supported by National Institutes of Health Grants R35 GM118091 and R01 AI119178. The NMR experiments were performed at the National High Magnetic Field Laboratory, funded by the National Science Foundation Division of Materials Research (DMR-1644779) and the State of Florida. We would like to acknowledge Steven Weigand of the DND-CAT staff for helping to collect SAXS data at the Advanced Photon Source of the National Argonne Laboratory.

